# Self-driving development of perfusion processes for monoclonal antibody production

**DOI:** 10.1101/2024.09.03.610922

**Authors:** Claudio Müller, Thomas Vuillemin, Chethana Janardhana Gadiyar, Jean-Marc Bielser, Jonathan Souquet, Alessandro Fagnani, Michael Sokolov, Moritz von Stosch, Fabian Feidl, Alessandro Butté, Mariano Nicolas Cruz Bournazou

## Abstract

The development of autonomous agents in bioporcess development is crucial for advancing biopharma innovation, as it can significantly reduce the time and resources required to transition from product to process. While robotics and machine learning have greatly accelerated drug discovery and initial screening, the later stages of development have primarily benefited from experimental automation, lacking advanced computational tools for experimental planning and execution. For example, in the development of new monoclonal antibodies, the search for optimal upstream conditions (such as feeding strategy, pH, temperature, and media composition) is often conducted using sophisticated high-throughput (HT) mini-bioreactor systems, while the integration of machine learning tools for experimental design and operation in these systems have not matured accordingly.

In this work, we introduce an integrated user-friendly software framework that combines a Bayesian experimental design algorithm, a cognitive digital twin of the cultivation system, and an advanced 24-parallel mini-bioreactor perfusion experimental setup. This results in an autonomous experimental machine capable of: (1) embedding existing process knowledge, (2) learning during experimentation, utilizing information from similar processes, (4) predicting future events, and (5) autonomously operating the parallel cultivation setup to achieve challenging objectives. As proof of concept, we present experimental results from 27-day-long cultivations operated by the autonomous software agent, which successfully achieved challenging goals such as increasing the viable cell volume (VCV) and maximizing the viability throughout the experiment.

## 1. Introduction

Competition in the biopharmaceutical industry has driven many advances in process development and clinical manufacturing for recombinant proteins. Today, R&D teams are challenged in early development phases to deliver a product within very short timelines. To achieve this, High Throughput (HT) devices are key to accelerate the development process of new molecules in the biopharmaceutical industry pipeline. Robotic platforms are used to increase experimental throughput and deliver the best production strategy through Quality by Design (QbD) approaches (Rouiller et al., 2012; Saleh et al., 2021). Robocolumns are for example used for downstream process (DSP) condition screening (Baumann et al., 2015). Microwell plates are used to cultivate cells during cell line development and upstream process (USP) development (Rouiller et al., 2016). Other advanced robotic systems enable to isolate cells in a single pen of 1.7 nL on a chip containing 1750 pens, the Beacon from Berkley Lights (Rienzo et al., 2021). Also, devices that run perfusion cell cultures at a scale of only 2 mL with fully automated fluid controls and on-line measurements (Mobius Breez, MilliporeSigma) are commercially available (Schwarz et al., 2023).

While in drug discovery (Carracedo-Reboredo et al., 2021; Dara et al., 2022) and at the initial screening stages, different machine learning tools are used for micro-well plates experiments (Vamathevan et al., 2019) the design of advanced cultivation strategies that include optimal media composition and optimal profiles of feeding, pH and temperature among others, require larger vessels as well as advanced monitoring, and control systems (O’Flaherty et al., 2020; Pogodaev et al., 2024). Additionally, with an increasing complexity on modern production processes as is continuous production (Fisher et al., 2019), experimental design and operation face new challenges. As stated in the thorough review by (Khuat et al., 2024), the current literature lacks a comprehensive solution for the autonomous operation of parallel cultivation systems with advanced control. Furthermore, experimental setups in high throughput that properly mimic industrial conditions are key to maximize speed and robustness during scaleup (Anane et al., 2018) and promote development following the QbD principles. In the field of cell culture, this has driven the development of highly sophisticated experimental systems, such as the commercially available family ambr250® robots (Sandner et al., 2019) Figure 1. This stage represents a significant bottleneck in development, as even with the presence of robotic systems, the design and execution of experiments still heavily rely on human intervention.

**Figure 1:**
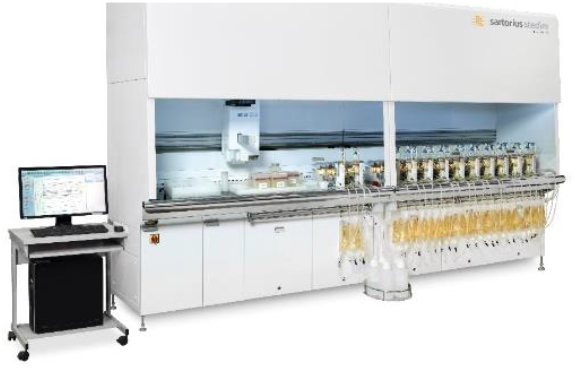
ambr®250 perfusion system

Emulating industrial process conditions is key to accelerating scaleup and minimizing the risk of failure. Nevertheless, performing proper scale-down experiments for cell cultures poses significant challenges (Karst et al., 2018). Such parallel cultivations demand a tight control over several critical factors such as agitation, aeration, feeding, pH and temperature among others. The complexity of the system is further increased with the incorporation of long-term membrane perfusion cultivations. This setup requires simultaneous control of additional media inlet flow, continuous harvest of liquid bulk to maintain constant working volume, cell retention within the reactor, and a bleed output for process purging. Despite the high level of automation in these systems, they still heavily rely on manual monitoring and operation. While low level controls (feedings, pH, temperature) are well implemented, computational tools that make important high-level decisions during operation are currently missing. Data acquisition also presents challenges, with parameters like pH, dissolved oxygen, and temperature being measured online, while metrics such as viable cell density, viability, and glucose concentration are assessed at-line. Furthermore, certain quality attributes may take weeks to quantify (Jang et al., 2009). All these aspects make it difficult for scientists to optimize the experiments while they are running. Also, the necessary computational tools to support operators and perform autonomous experiments are missing. This is in contrast with both, the ever-increasing number of computational tools developed and applied in biopharma in manufacturing and other development stages (Harrer et al., 2024), and the technologies already deployed in other fields, for example in material science and chemistry (Aspuru-Guzik, 2022).

Self-driving research is pushing the boundaries of digital discovery by giving full autonomy to “robot scientists” (Kramer et al., 2023). These systems can perform experiments with minimum human intervention, learn from their experimental results, and decide how to continue experimentation (Abolhasansofti and Kumacheva, 2023). While self-driving clearly suggests an association with mobile systems (Burger et al., 2020), the use cases of self-driving laboratories rarely deal with dynamic systems and the challenges hereby involved (Duong-Trung et al., 2023). This has evidently been no limitation to achieving incredible results in chemistry, catalysis, material science, and biology (Boiko et al., 2023) but process dynamics become increasingly relevant as we approach industrial scale production (Haringa et al., 2018; Villiger et al., 2018). To develop self-driving systems for efficient bioprocess development and scale-up, it is essential to incorporate process dynamics and control within the framework (Chopda et al., 2022). We need hence a digital twin (DT) of the dynamic cultivation process to ensure an optimal design of the experimental campaign and its proper operation by automated systems (Narayanan et al., 2020). Furthermore, due to system uncertainty and discovery related nature of experiments in R&D, the DT requires important cognitive properties (Lu et al., 2022). The capability to use existing knowledge and learn from the experiment that is being operated (Cai et al., 2020) is essential to ensure the fastest development possible and efficient process control of well-designed experiments (Kim et al., 2023).

These solutions need, furthermore, to be user friendly to enable its use by experts in experimentation of mammalian processes. Furthermore, the implementation of robust tailor-made computational pipelines and development of corresponding computational workflows that can automatically handle and store all the data and metadata generated before, during, and after conclusion of an experiment, play an essential role (Bai et al., 2022; Mione et al., 2024). Unfortunately, speaking of miniaturized bioreactor cultivations, while the experimental tasks have been automated to great extent and the throughput increased significantly in the last decade, less advances have been achieved in embedding automated computational workflows for data management and decision making.

As a result, while modern experimental robots can perform very complex tasks automatically for long periods of time, experts’ intervention is still regularly required to analyze the data and to operate the parallel cultivations. This has to be considered in the context of the large experimental time spans (up to months of cultivation) and the high costs of the experiments reaching 50 k€ in some cases. Finally, the high degree of automation of the experimental setup necessitates an integrated decision-making agent, to minimize the human involvement in these lengthy and time-consuming experiments. Algorithms that make the best decisions to operate several cultivation processes in parallel maximizing the knowledge derived from such experiments, are pivotal for accelerating development.

In an effort to tackle these issues and significantly reduce time and costs in the development of monoclonal antibody manufacturing strategies, we present a user-friendly software solution that enables the autonomous operation of the parallel bioreactor system with perfusion membrane modules. The key features are:

1. Hybrid modelling formulation for flexible and robust process
2. Online model re-training to enable real-time learning
3. A toolset to transfer learnings from past projects
4. Predictive process monitoring and notifications autonomous feed-back experimental operation minimizing the human in the loop

The results demonstrate the capabilities of step-wise Gaussian process models (SW-GP) to learn from the data, to transfer learnings between different cell lines, and to support optimal experimental design with Bayesian methods. They also show that the optimal operation can robustly fulfill challenging tasks throughout longer periods of time and maintain critical conditions as is cell viability within specification.

## 2. Material and Methods

Short introduction to the experimental setup and operation

All experiments presented in this study were performed in an ambr®250 system with 24 bioreactors vessels of 250mL volume and a membrane module to enable perfusion cultivations Figure 1

### 2.1. Cell expansion

Cells were thawed and diluted at 0.30 10^6^ cells/mL in 50 mL SpinTube (TPP, Trasadingen, Switzerland). Cells were diluted every 2 or 3 days to 0.30 10^6^ cells/mL or 0.20 10^6^ cells/mL respectively and using proprietary expansion medium. Cell culture volumes were increased from SpinTubes to 2L wavebags (Sartorius, Goettingen, Germany). After two weeks of expansion, cells were transferred to inoculate the perfusion cell cultures on the AMBR250 (Sartorius, Goettingen, Germany).

### 2.2. Perfusion process

ambr®250 perfusion bioreactors were inoculated with cells at the maximum concentration available in 213 mL of final volume. Cells were cultivated in a proprietary chemical defined medium. During the growth phase, the perfusion rate was increased from 0 to 1.3 d-1 until reaching the target viable cell volume (VCV) which was calculated according to the following equation,

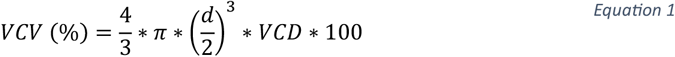

where d is the average cell diameter and VCD the viable cell density (10^6^ cells/mL).

The perfusion volume was maintained constant at 213 mL throughout the duration of the run. Once the VCV target was reached, the culture was considered to be in a state of control and the given setpoints for the input parameters were followed (see Table 1 and Table 2). The bleeding was triggered if the value of VCV was higher than the setpoint. Sample analysis of the bioreactors was performed daily using a Flex2 (Nova Biomedical, Waltham, MA, USA) to count the cells, measure cell viability (Via), average cell diameter (Diam), quantify glucose (Glc), glutamine (Gln), glutamate (Glu), lactate (Lac), ammonium (Amm) concentration, pH, pO_2_, pCO_2_ and osmolality. Samples for off-line analysis were taken to quantify monoclonal antibody (mAb) titer, amino acid concentrations and quality attributes (glycans, oxidation and deamidation) of the mAb produced. A glucose bolus was performed if the glucose concentration was lower than 2.00 g/L after the sampling. The feed volume was calculated to replenish glucose to a concentration of 5.00 g/L.

**Table 1:** Steady state experimental plan to build a design space where the digital twin will build and update basal models

| Parameter | -1 | 0 | 1 |
| --- | --- | --- | --- |
| Perfusion rate (Vessel Volume per Day (VVD, d<sup>-1</sup>)) | 0.5 | 1.25 | 2 |
| Temperature shift after reaching VCV target (°C) | 34 | 35.25 | 36.5 |
| Stir speed (rpm) | 700 | 1050 | 1400 |
| Pyruvate feed flowrate (g.L<sup>-1</sup>.d<sup>-1</sup>) | 0 | 1 | 2 |

**Table 2:** Clone tested, control variables and output variables for each bioreactor. Variables to be controlled could change in the tested ranges of Table 1 except for stir speed which could evolve between 700 to 1050 rpm as higher rpm was deleterious for cell viability.

| Condition | Clone | Variable to be controlled | What to optimize |
| --- | --- | --- | --- |
| 1 | Clone A | VVD, Temperature, stir speed, pyruvate flow | Target 30 % VCV |
| 2 | Clone A | Pyruvate flow | Target 1mM NH<sub>4</sub><sup>+</sup> |
| 3 | Clone B | Pyruvate flow | Target 1mM NH<sub>4</sub><sup>+</sup> |
| 4 | Clone B | VVD, Temperature, stir speed, pyruvate flow | Target 98 % viability |
| 5 | Clone B | VVD, Temperature, stir speed, pyruvate flow | Target 98 % viability |
| 6 | Clone B | Pyruvate flow | Target 1mM NH<sub>4</sub><sup>+</sup> |

### 2.3. Training experiment

A perfusion experiment consisting of 24 runs with a fixed experimental design according to Table 1 was performed with one cell line CHO-K1 Clone A. The purpose of this run was to gather data with sufficient variability for the initial model training using standard process conditions. This experiment is referred to as the Training experiment.

Once the VCV target was reached, all parameters gathered in Table 1 were changed according to the design of experiment.

### 2.4. Use case experiment

The second perfusion experiment (Use case experiment) was performed with two cell lines: Clone A (same clone as Training experiment) and Clone B (new clone). Once the state of control was reached, inputs suggested as setpoint by the model aiming to comply with the constraints and reach the objectives given to each bioreactor (Table 2). All other set-points were kept as in the previous experiment.

## 3. Computational Methodology

In this section, we first discuss the communication with the experimental system using a Structured Query Language (SQL) database. We then shortly discuss the computational framework based on hybrid modelling and Bayesian optimization to finally give a detailed description of the experimental setup.

### 3.1. Gateway connection and data storage

As more laboratories adopt HT experimental technologies and the amount of data generated per time increases, systems for HT data treatment and storage are imminent to assure the traceability and provenance of the experiments as well as to keep up with the data generation speed.

The communication between the cultivation system, here the ambr250, and the software is based on a ClientServer Architecture, over TCP/IP standards developed together Institutsleiter Elektrotechnik, Hochschule Luzern T&A.

In the current framework, the bioreactor system interfaces with an OPC UA Server (KEPServerEX), from which data transmissions to an OPC client over TCP/IP, to a MySQL Database, allowing to collect and store the data from the experimental system. The software interfaces with the SQL client, allowing to query the process data stored in the MySQL Database. Most importantly, the software is also able to send experimental set-points to the OPC client, which are then enacted in the bioreactors with low level control in real time. Additionally, all metadata is stored in an Excel macros file (.xlsm) and the actions performed by the robots is recorded in (.csv) files and stored to ensure full traceability of the robotic actions during experimentation (Mione et al., 2024). The communication architecture is shown schematically in Figure 2.

**Figure 2.**
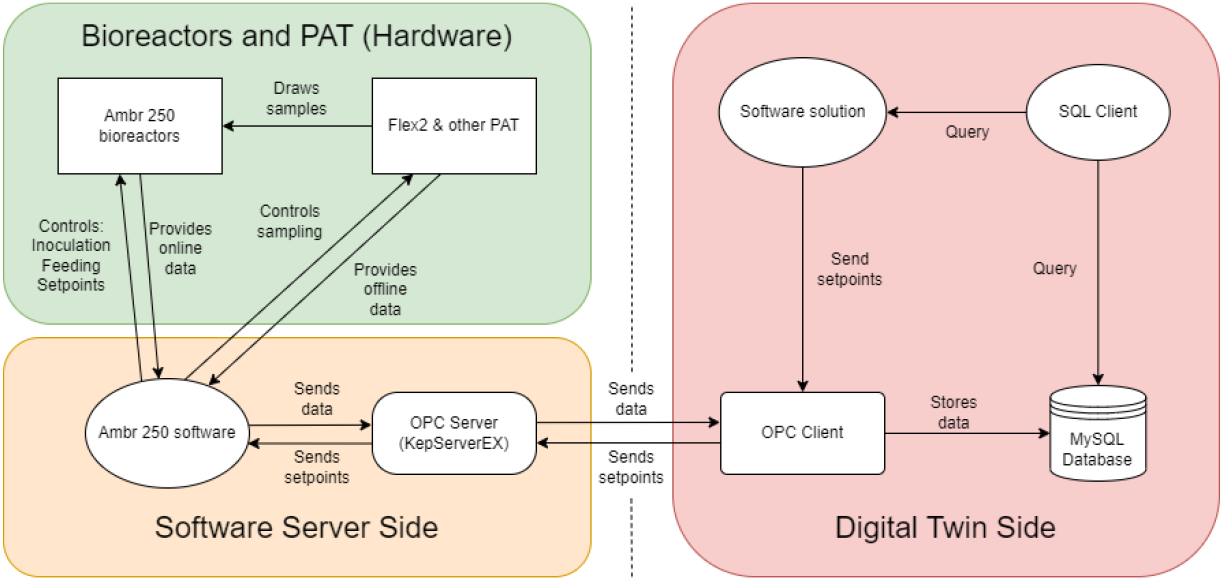
Overview of the interconnectivity of the DT software solution, the OPC and SQL client, as well as the ambr®250 server.

### 3.2. Hybrid gaussian process model

The mathematical model used to describe the evolution of the bioreactors during experimentation needs to consider a system with a vast number of cellular biochemical reactions that take place in bioprocess cultivations (González-Hernández and Perré, 2024). In a bioreactor, the chain of reactions that take place in the cells for metabolite secretion and/or antibody production might be unknown. The existing models might not capture all the factors affecting the reactions, for exeample pH and temperature dependency on shear rates. The presence of impurities in the media can also affect cellular reactions and are very difficult to detect. Due to these reasons, purely mechanistic model might not be able to accurately predict process behavior in a bioreactor without re-calibration (Cardillo et al., 2021). Hybrid modelling is an alternative approach, which combines the advantages of mechanistic and data-driven models to better describe complex systems (Narayanan et al., 2019).

For our current purposes, we use a hybrid model formulation based on basic mass balance equations represented by a system of ordinary differential equations (ODE) combined with Gaussian Process (GP) models as the data-driven counterpart of the hybrid model (Azevedo et al., 2019; Mahanty, 2023). GPs use a measure of similarity (kernel function) between points in the training data to predict the distribution of a value for an unseen datapoint. With the kernel function being usually non-linear, such an algorithm is capable to reproduce highly non-linear and complex behavior. The major advantage being the estimation of both, the predicted value and its associated uncertainty (Kocijan et al., 2004).

As reported in literature, Gaussian Process State Spaces Models (GPSSM) (Umlauft et al., 2017), that aim to describe the state space with GPs, have shown to also represent mammalian cultivations well (Cruz Bournazou et al., 2022). This formulation allows to use GPs to describe nonlinear dynamic processes with limited knowledge of the phenotype and growth dynamics of the cultivation (Hutter et al., 2021). The advantages of GPSSMs have been documented in several applications with small to mid-size data sets with noise that is close to Gaussian (Kocijan et al., 2005).

Given the slow dynamics of mammalian processes, results show that discretion with a time step of up to 24 hours is sufficient to properly mimic the evolution of the process over time. Based on this assumption, we can significantly reduce the computation burden implementing a SW-GP as described here.

The time discretized ODE system is formulated as:

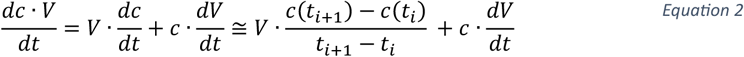

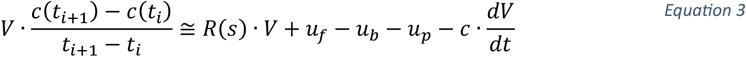

where *c* is a vector of concentrations (e.g., Viable Cell Density (VCD), glutamine, glucose, lactate, ammonium, titer); *s* a vector of process states (i.e., all time dependent variables, including the concentrations *c*); *V* is the culture volume; *t* is a vector with time stamps *t*_*i*_; *u*_*f*_, *u*_*b*_, and *u*_*p*_ vectors of mass feed rates (nonzero only for compounds that are fed), mass bleeding rates, and mass perfusion rates, *c*(*t*_*i*_) is the concentrations measured at the step *i*, and *c*(*t*_*i*+1_) the concentration measured at the following step, *i* + 1. For the sake of simplicity, in Equation 3 all quantities on the right-hand side are evaluated at time *t*_*i*_.

Note that the change in bioreactor volume over time, *dV*/*dt* can be computed explicitly. Using the discrete hybrid model, called SW-GP from now on, it is possible to calculate the term *R*(*s*(*t*_*i*_)) at every time step *t*_*i*_, as all other terms are measured, by rearranging Equation 3:

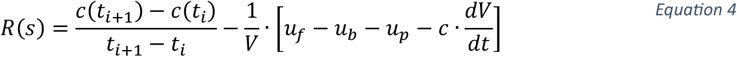

We now can explicitly compute the discrete rate of production for every species at the step *i*, and relate such term to the state of the process measured at the step *i, i*.*e*., *s*(*t*_*i*_). In the SW-GP formulation, this is done using a GP model, *i*.*e*.: *R*(*s*) *≈ GP*(*s*). The inputs to the GP are the process states *s* measured at every time *t*_*i*_, while the outputs are the corresponding discrete rates of production, *R*.

The formulation of the discrete hybrid model of Eq. 2 allows to learn the process evolution in time for each variable in a stepwise fashion. Once the initial condition *s*(*t*_0_) are defined, where *t*_0_ = 0, it is possible to compute the rate of production for each state variable in the vector *c*(*t*_*i*_) corresponding to the step (*t*_*i*+1_ − *t*_*i*_) and, using the discretized mass balance above, compute e.g. the value *c*(*t*_1_) at time *t*_1_. At this point, the state of the process at the new time, *s*(*t*_1_), is defined and the procedure can be repeated for all steps.

The SW-GP model is trained as an ensemble of smaller Gaussian Process models. The model consists of many sub-models, which are individually trained on randomly sampled subsets of the full training data. For each prediction, the sub-models individually predict the process dynamics, and these predictions are used to calculate confidence intervals. Specifically, the 10^th^ to 90^th^ percentiles of the aggregated predictions provide the 80% confidence interval, allowing for uncertainty estimates. The median prediction is given by the 50^th^ percentile.

The relative root mean squared error (rRMSE) is used for both, training and evaluation of the SW-GP. The model was evaluated considering its capability to predict the next three daily timesteps, since this is the time horizon considered during operation.

The overall error of the model for a variable *x* is given by,

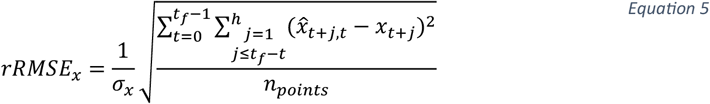

where *t*_*f*_ is the process duration in days, *t* the current process day from which the model predicts, *h* the prediction horizon of interest (here three time steps), 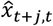 the prediction of variable *x* from day *t* to day *t*+*j, x*_*t*+*j*_ the future observed value on day *t*+*j, σ*_*x*_ the standard deviation of the respective X variable and *n*_p_ the total number of points to be averaged over, given by,

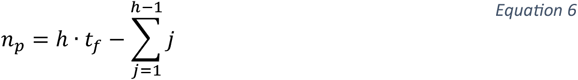

### 3.2. Online retraining of hybrid models (example with in-silico data)

Most applications for model-based operation of robotic experimental facilities work either offline or decouple the learning phase from the experimental plan. Yet, a Cognitive Digital Twin, with the ability to learn on the fly during experimentation is needed to ensure an efficient experimental campaign of new clones and/or processes. Standard methods for adaptive control, for example, still rely on mechanistic models and reliable estimates for the initial parameter values (Krausch et al., 2022). Additionally, the transfer of learnings from previous processes to new ones is essential to ensure the use of all available knowledge and a rapid development of new products (Hutter et al., 2021). To tackle this, frameworks that are capable of both learning form experiments and transferring previous learnings are needed. Unfortunately, the tools that have been developed up to now for biopharma are very specific and require expertise in modelling, optimization, and programming to be adapted to new scenarios (Narayanan et al., 2020). Today, R&D in biopharma is missing the necessary toolset to fully exploit the potential of parallel mini-bioreactor systems and speed up the generation of knowledge to reach truly accelerated innovation.

The main challenges related to the design and operation of informative experiments in R&D can be strongly alleviated by 1. the transfer of existing knowledge from similar processes and 2. a frequent re-calibration of the models as data is being recorded, to maximize the efficiency of long experimental campaigns (several weeks) of experimentation. To show the added benefits of online retraining a model during an ongoing experiment, in-silico datasets for two different clones (clone X and clone Y), corresponding to a dataset size with 24 run each, were generated using experts’ knowledge of the real process. The data was generated using a mechanistic model that describes fed-batch cultivation of mammalian processes with lactate consumption (clone X) and without lactate consumption (clone Y). A fed-batch process over 14 days of cultivation with similar conditions to the use case (see M&M) was considered to generate the insilico data. The state variables are VCD, titer, glucose (Glc), glutamine (Gln), ammonium (Amm), lactate (Lac), dead cell density (DCD) and lysed cells (Lysed). The input variables that were controlled are the stirring rate set-point, the DO set-point, initial conditions of glutamine, glucose and VCD, and the feeding volumes. The hybrid model was initially trained using only data from clone X with 24 runs. The experimental design was simulated with each insilico data point treated as a daily measurement. On each timestep, the model is re-trained using the existing insilico data, prediction calculated for the next three days. In the first, the initial model of clone A is used, without adaptations. In the second, the new process data of the clone B runs is added to the training data and the model is retrained (see Figure 3).

**Figure 3.**
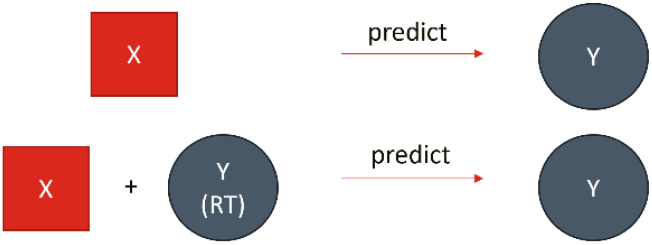
Schematic representation of model comparison approach. In the first case (top row), the model is only trained on clone A and used directly to predict runs of clone B. In the second approach, the model is initially trained on clone X, but then continuously retrained with current available clone Y data during the process (RT).

Prediction errors were then computed for a 3-day horizon considering the operation scenario. The results of the above comparison are shown in Figure 4, where the rRMSE was determined as described previously. The results depicted confirm that, as expected, the model predictions are more accurate in the case that states and model parameters are re-trained continuously. This trend is indeed exemplified for processes which run for long durations, which is typically the use case and purpose in perfusion processes.

**Figure 4.**
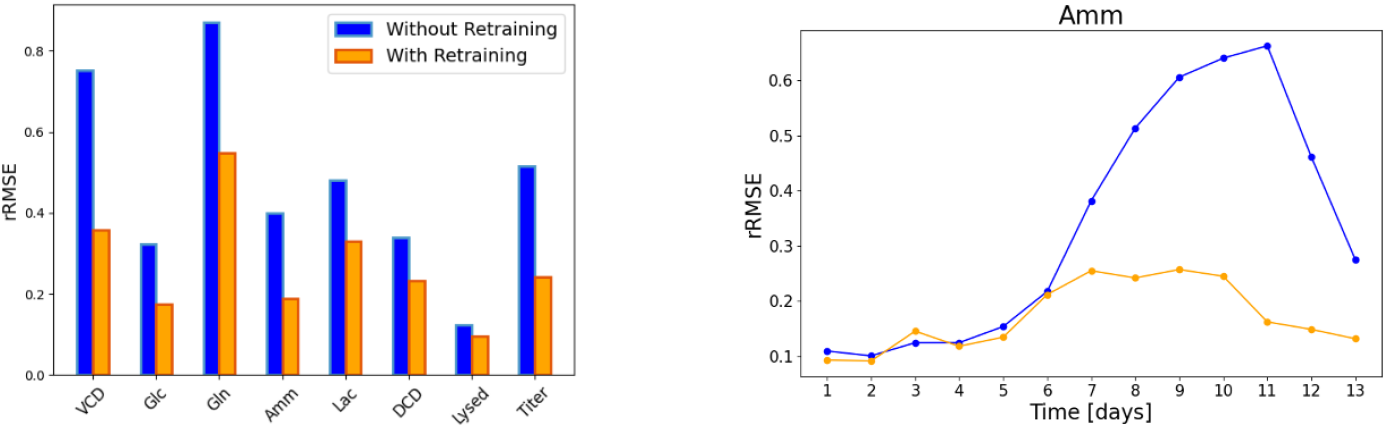
Left: Comparison of relative RMSE results when the model is not being retrained (blue) and when it is retrained (orange). Predictions are evaluated over a 3-day horizon. Right: Relative RMSE of Ammonia samples as a function of the process day. Model errors were averaged over all 24 runs in the simulated validation campaign. For the error calculation, only 3-day ahead predictions were considered until day 11.

### 3.4. Optimization framework

Designing experimental campaigns involving multiple parallel runs, extended cultivation durations, and numerous input variables to optimize poses a significant challenge for Optimal Experimental Design (OED). OED and optimal experimental operation problems have been tackled for many decades by the Process Systems Engineering (PSE) community (Franceschini and Macchietto, 2008) and has also rapidly gained popularity in the machine learning community (Rainforth et al., 2023). There are also specifically tailored for bioprocess cultivations (Cruz Bournazou et al., 2017; Kim et al., 2021; Martínez et al., 2021).

A very promising approach with the advent of machine learning is to use Bayesian Optimization to treat the problem as a Bayesian Experimental Design (BED) task (Rainforth et al., 2023) also with several applications for parallel systems (González and Zavala, 2022; Luna et al., 2025).

The optimization framework based on Bayesian Optimization (BO) searches for the best experimental strategy within the parallel bioreactor system, aiming to maximize the probability that the clone and operating conditions are found. The BO framework searches operating conditions (in particular, temperature, agitation rate, vessel volume per day (VVD) and pyruvate additions) using the SW-GP model described in 3.2 to compute the acquisition function.

The objective function is constructed by defining a target value for the variable to optimize and by considering only the model predictions of the next 3 daily timesteps. The predictions utilized to evaluate the performance of the process at the given parameter values aiming to find the optimal inputs to bring the process closest to the defined target.

Mathematically, this is described as,

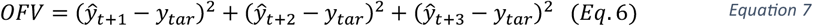

where *OFV* is the objective function value to be minimized, 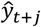 is the predicted value of the process variable *y* at *j* days in the future and *y*_*tar*_ is the desired target value of *y*. The inputs that were chosen to be varied were the set-points of temperature, agitation rate, VVD (perfusion rate) and pyruvate addition.

### 3.5. User interface

Finally, in order to enable a human centric digitalization of the development process in compliance with bioprocessing 5.0 (Xu et al., 2021), a user interface was developed together with expert users and technical operators.

The software solution was equipped with multiple features, which are briefly described below, where some are supplemented with a screenshot of the respective interface:

- Main Overview (Figure 5 top left): Visualization of all measured data for all ongoing bioreactor runs, as well as past experiments. The user may choose any process variable, highlight individual runs or remove them from the visualization.
- Comparison: Multiple univariate data visualizations in one interface, allowing fast comparison across variables and ongoing runs.
- Model evaluation (Figure 5 top right): Overview of all past model predictions for a single run in focus. Includes observed vs. predicted for a given process day as well as rRMSE metrics.
- PCA (Figure 5 bottom left): Multivariate data analysis of all ongoing runs, allowing to identify key correlations between variables, as well as detecting outliers.
- Notifications: The user may set up alarms regarding violation of critical limits of important process variables. The model predicts if constraints will be violated in the future and informs the user on a daily basis if this will be the case.
- OPC: In this section the current connection status is shown and can be queried.
- Optimizer (Figure 5 bottom right): Here the objective functions for optimization are defined on an individual basis for each bioreactor. The control parameters (such as temperature, perfusion rate, agitation rate and pyruvate addition) are chosen, as well as the lower and upper bounds for search space definition. For the target variables a target value is defined, or they are simply maximized or minimized.
- Log: Record of all actions taken by the software, such as data queries, written set-points and connectivity status.
- Data: Tabular visualization of all process data of ongoing or past runs.

**Figure 5.**
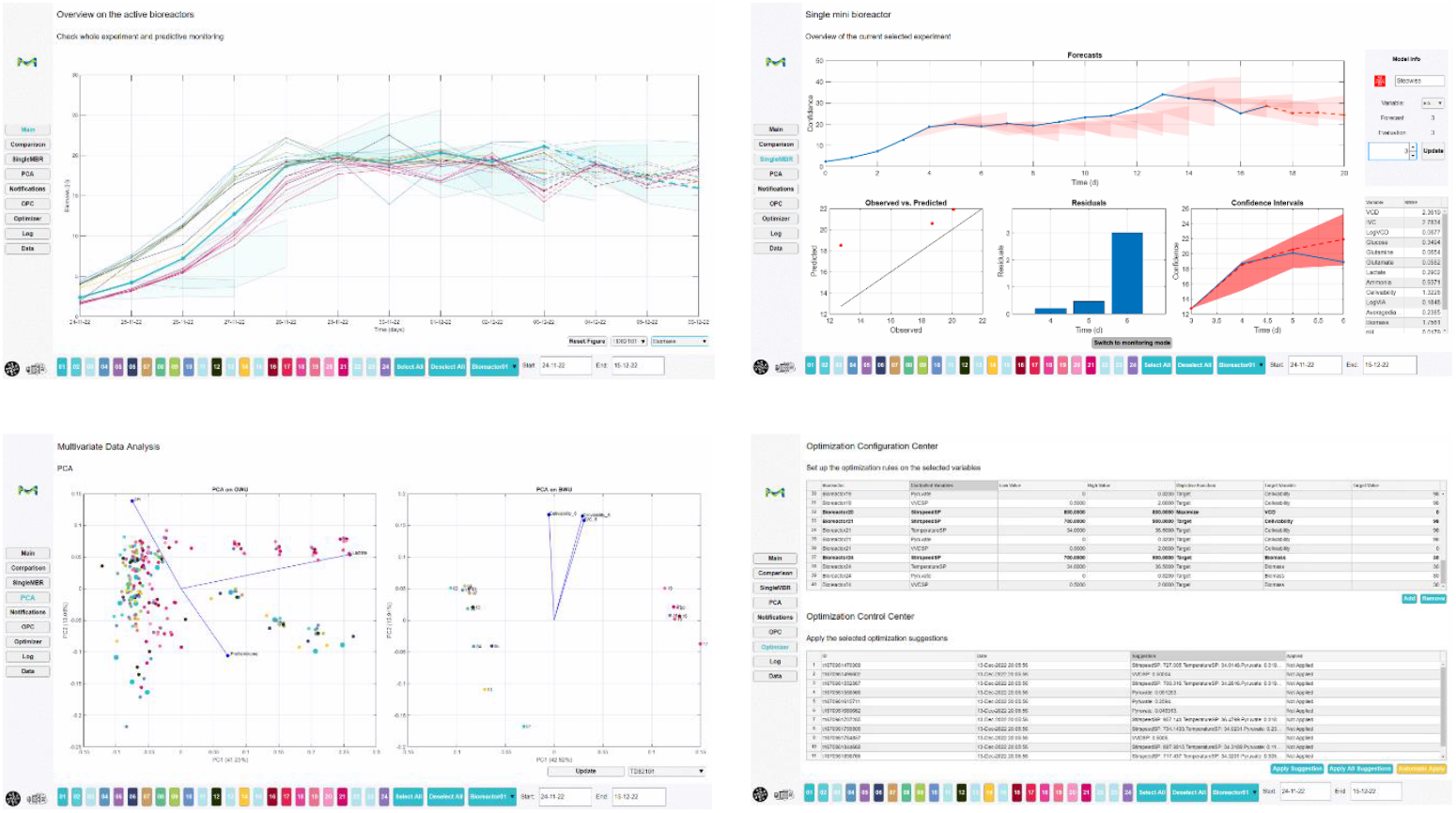
Screenshots of the user interface developed for easy use and implementation of the software: (top left) main overview of active bioreactors, (top right) Model evaluation for a single run. Past forecast’s prediction intervals are shown in the top plot, as well as current predictions, (bottom left) PCA plots of the variable- or observation wise unfolded (OWU) matrix on the left and batch-wise unfolded (BWU) matrix on the right, (bottom right) optimization configuration center. For each bioreactor, the control variables are configured in terms of lower and upper bounds to define the optimization search space.

## 4. Results

### 4.1. Experiment for initial model calibration

Specific experiments were tailored to collect data, as described in Section 3.3 and were used to train a hybrid model as explained in Section 2.2. The model output accuraccy was assessed by training models in leave-one-out validation. The model errors were determined as described in Section 2.5. Figure 6 shows the rRMSE obtained for the trained model against the validation data sets. The left bar plot shows the average rRMSE over all runs in testing over a 3-day horizon. For all variables the predictions have a rRMSE below 0.5 in the 3-day prediction horizon considered, making it suitable for control. On the right-hand side the rRMSE is displayed at the resolution of the runs for each variable. There are no runs (rows), where all variables are predicted with significantly higher rRMSE than the overall results, as can be seen by the intensity of the blue shadings. Therefore the heat map is indicating that the model can generalize well on the entire design space.

**Figure 6.**
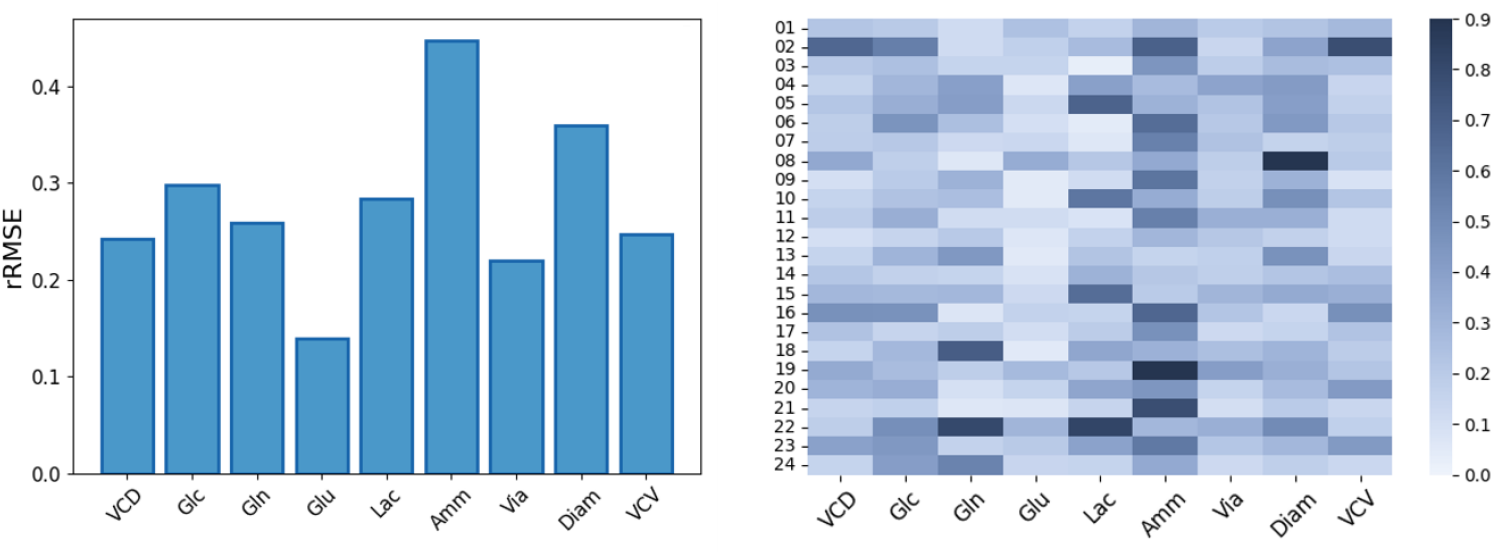
Left: rRMSE results averaged over all runs in testing (leave-one out) over a 3-day prediction horizon. The relative RMSE is well below 0.5 for all variables indicating good model performance. Right: rRMSE heat map showing the errors for each run (row) and variable (column).

In order to better visualize the model prediction capability, a representative run in terms of model performance (labeled as “01” in Figure 6, right-hand side) was picked for the purpose of illustration. Figure 7 shows the median model prediction and the prediction interval for the following 3 days on each day of operation. We can clearly see that the model predicts the trends of the process accurately and in general the observed process data is within the prediction interval of previous predictions.

**Figure 7.**
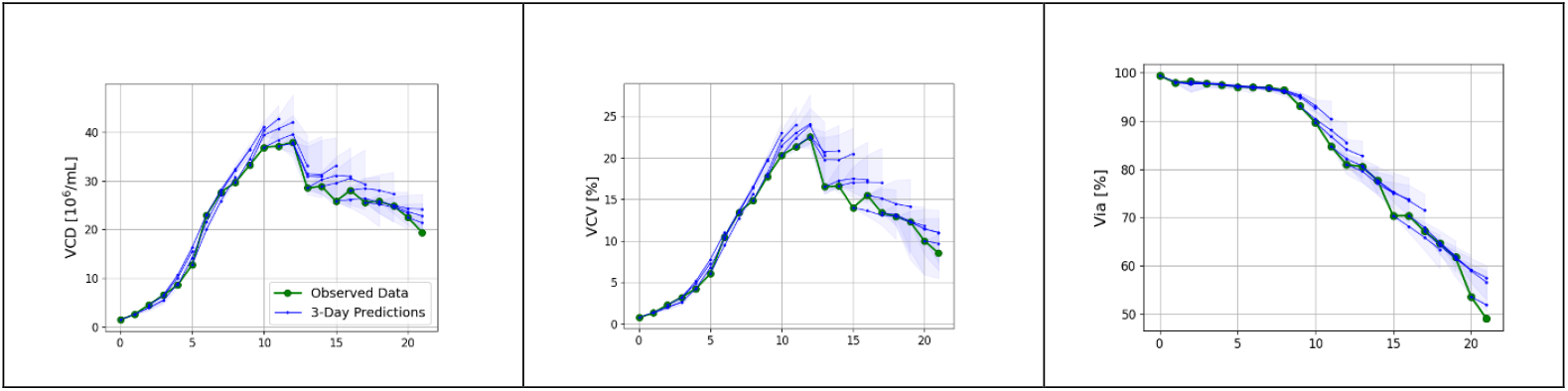
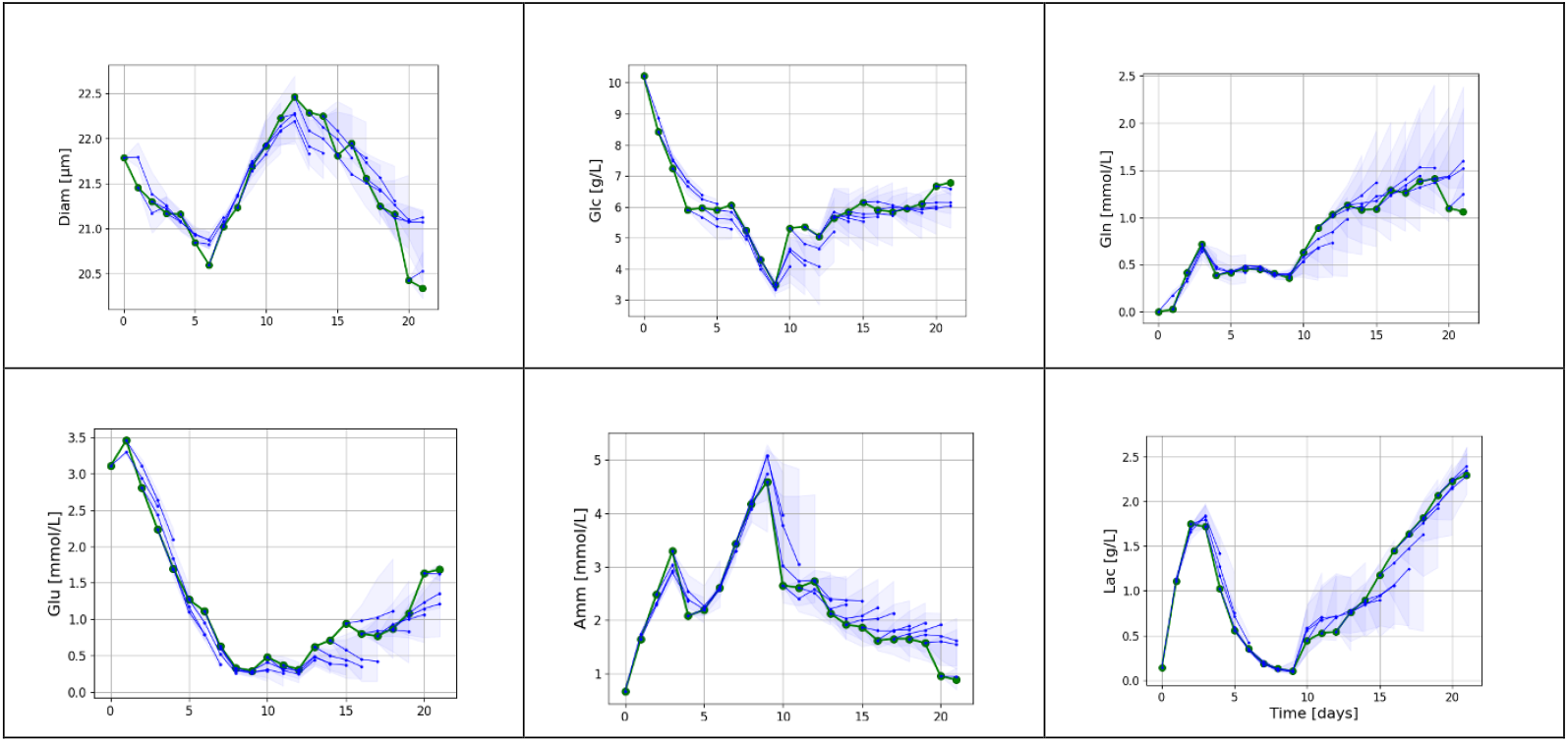
Predictions for a run in the Training experiment in testing. The model was trained on the remaining 23 runs. The green line shows the measured experimental data, the solid blue lines are the 3-day median predictions of the model, starting on each respective day. The shaded polygons represent the prediction intervals. In the evaluated 3-day horizon, the model predicts the process trends with sufficient accuracy for software operation.

The results demonstrate that the initial model can learn the process behavior and can make meaningful predictions across different input process conditions.

### 4.2. Demonstration of software performance with use case experiments

Three different use cases were selected to target process indicators such as VCV, Viability and ammonium, in order to evaluate the online control of the bioreactors by software solution as well as it’s optimization and retraining capability. The control variables selected for all use cases are: perfusion rate, temperature, agitation rate, and pyruvate addition. Each day, the software would read the data provided by the robotic platform and re-compute the optimal control variable setpoints to reach the defined target for each use case.

The different use cases demonstrate that a training experiment which covers the expected variability for given process parameters will help the hybrid model to learn how control variables impact process indicators. This knowledge from the model can then be used to operate the process once the desired state of control is reached. Deviations from the state of control can be predicted and avoided with a correct response from the system. This strategy is applicable for process development but could also be useful in manufacturing. Debates on how to validate such models for GMP applications should help the biomanufacturing community to exploit such capabilities to their full potential.

#### 4.2.1. Use case 1: increase of the Viable Cell Volume

The objective set to the software agent for this use case was to increase VCV to a target value = 30%. To achieve this, the setpoints of all four control inputs were varied every 24 hours as shown in Table 2, Condition 1.

Figure 8, left shows the VCV evolution during the process for the bioreactor with software operation (green), 3 bioreactors of the same clone without software operation (blue). Perfusion rate (Perfusion) is the control variable that has the biggest impact on the target and is shown in light red in the large plot. The smaller subfigures at the top of the figure show the set points suggested by the optimizer for the other control variables, namely temperature shift (Temp), agitation rate (Stir) and pyruvate addition flowrate (Pyr) respectively.

**Figure 8.**
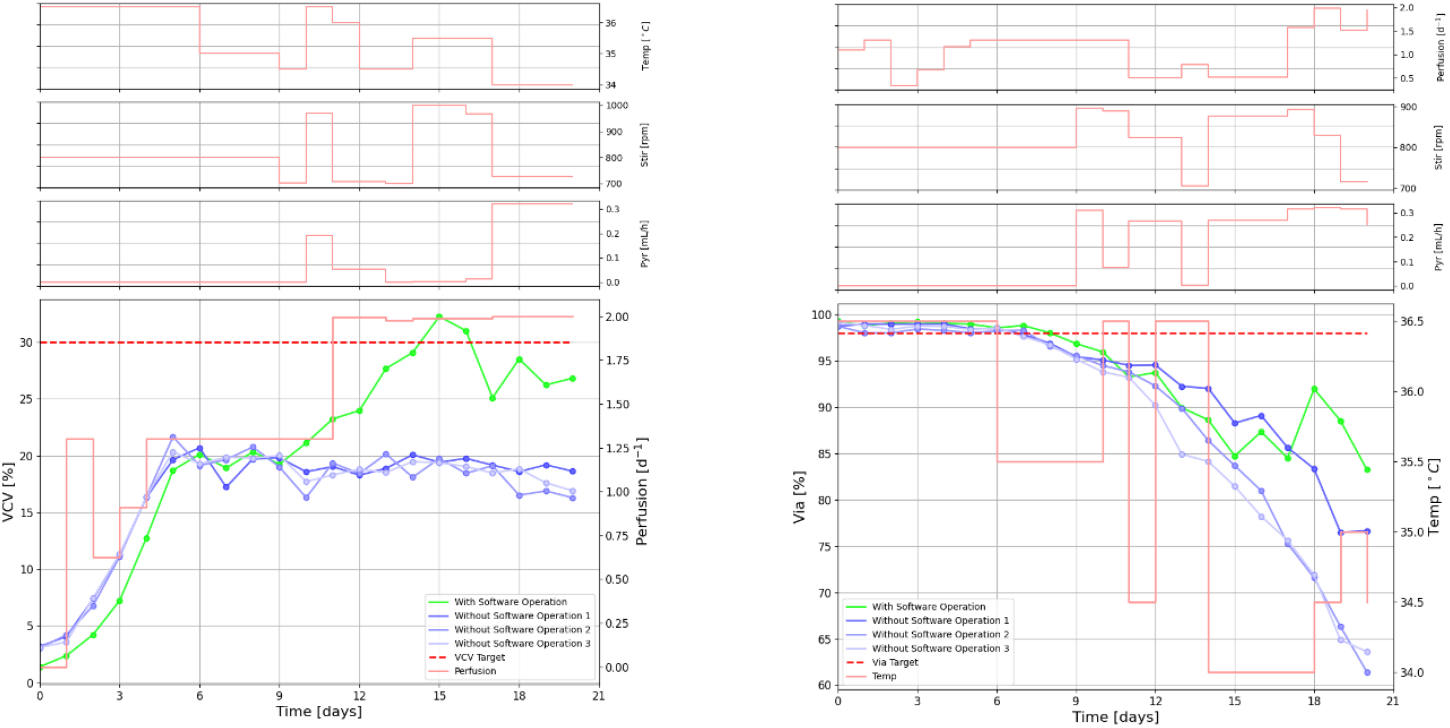
Left: Condition 1 results: VCV use case with VCV Target = 30% (red dashed line). The observed VCV values of the bioreactor with software operation are shown in green, the observed values of three bioreactors without software operation are shown in blue. The optimizer suggested perfusion rate is plotted on the secondary axis. All control parameter profiles are shown in light red. Right: Condition 4 results: viability (Via) use case with Via Target = 98% (red dashed line). The observed viability values of the bioreactor with software operation are shown in green, three bioreactors without software operation are shown in blue. The optimizer suggested temperature rate is plotted on the secondary axis. All control parameter profiles are shown in light red.

The optimizer was making suggestions from day 9 onwards. For this use case the VCV target value of 30% is significantly higher than the value of VCV in standard experiments (average VCV after state of control < 20%). The software was able to suggest conditions that significantly increased VCV reaching the target in contrast to the benchmark bioreactors.

The software identified perfusion rate as the most significant factor for controlling VCV. Higher perfusion rate will lead to faster replenishment of the perfusion media, thereby allowing faster cell growth. This is a well-known phenomenon among process scientists, which was inherently learned by the hybrid model using the training experiment data as well as the constant retraining of the model.

#### 4.2.2. Use case 2: Cell Viability

The aim of this use case was to maintain the cell viability = 98% of the clone B by varying all the control parameters as shown in Table 2, Condition 4. Since there exists no previous data on this clone, in the Training experiment, the capability to reach this objective demonstrates the transfer learning capabilities of the approach. Figure 8, right shows the cell viability evolution during the process for the bioreactor operated by the software (green), 3 bioreactors of the same clone without software operation (blue). In this use case, temperature is identified as the control variable which had the highest impact on the target and the set points suggested by the software solution are shown in red in the large plot. The daily optimized set points for the other control variables are plotted at the top of the Figure. The results of an additional viability use case are shown in Figure A2 in the appendix.

Similar to the previous use case, the online control of the software solution was started after day 9. The variations in the optimized set points, especially temperature and VVD were quite large until day 14 in comparison to the VCV use case. However, the response of the bioreactor to these changes was used to train the model. The retraining of the model with the new observed data led to an improvement in prediction of cell viability which in turn aids the optimization of control variables. This can be inferred from the increase in cell viability observed after day 15. The software solution suggested lower temperature set points as well as higher perfusion rate towards the end of the process. These conditions led to higher cell viability in the bioreactor in comparison to the bioreactors without software control. Apparently, the ability of the model to understand the impact of the process parameters on viability for the new clone at day 9 was not sufficient. Only after retraining with the data of the first half of the validation run, the optimizer was able to learn these relationships and re-optimize the set points accordingly.

#### 4.2.3. Ammonium use case

It has been suggested in literature that sodium pyruvate addition in a perfusion cell culture can be used to stabilize ammonium concentration (Caso et al., 2022; Romann et al., 2024). Pyruvate is the metabolic entry point for the Krebs cycle. If the glycolysis pathway is saturated and pyruvate becomes limiting, cell will use other sources (amino acids) to feed this cycle, and this can result in ammonium accumulation. Hence, the aim of this use case is to control ammonium concentration to a value = 1.0 mM by using pyruvate addition as the only control parameter. This is illustrated in Table 2 (Condition 2).

As depicted in the Appendix in Figure A3, the ammonium evolution during the process for the bioreactor with software operation (green), 3 bioreactors of same clone without software operation (blue) and model suggested pyruvate addition set points (red). We can see an increase in the ammonium concentration up to 4 mM until day 9 when there is no addition of pyruvate feed. The optimization results suggested the addition of pyruvate to reach the ammonium target, which led to a reduction in the ammonium value until day 13.

Manual injection of ammonium was performed on day 13 to confirm the response of the model to high values of ammonium in bioreactor, leading to the observed peak. Once the concentration of ammonium increased above 7 mM, the software responded by increasing the pyruvate addition set point, as it had learned that the addition of pyruvate decreases ammonium.

The results of two additional ammonium use cases, with identical targets and manual injections of ammonium, are shown in the appendix, Figure A1 and Figure A3.

## 5. Discussion

Novel robotic systems for HT experimentation at every lab scale have drastically changed biotechnology laboratories. The large number of runs that can operate in parallel, the highly complex tasks that robots can perform, the associated process analytical tools and the addition of mobile assistants that overcome spatial limitations, are just some of the many advantages. On the other hand, this significantly increases the number of high-tech experimental devices that rely on skilled personnel and on a robust digital infrastructure to enable communication, control, and orchestration. Some examples are data management tools, IT systems, optimization algorithms (for design or scheduling), monitoring and control tools.

As demonstrated in this study, the combination of computational tools with existing robotic experimental setups can drastically increase the throughput and efficiency of laboratories. The automated experimental design and execution of parallel perfusion experiments tackles a major bottleneck in bioprocess development. Moreover, the ability to continuously learn and adapt the experimental design during cultivation allows for real-time improvements as insights into the process are acquired. Especially for development of perfusion experiments that are expected to run for long periods of time, a repeated improvement of the experimental design is essential to maximize efficient use of time and resources.

The use cases presented in this work show that the software fulfills the 5 main requirements for self-driving laboratories for bioprocess development, namely:

- **Hybrid modelling formulation for flexible and robust process:** A robust model that allows a proper description of the time evolution of performance even for new clones is the backbone of an efficient DT of the process. The implemented hybrid model based on SW-GP was shown to describe the process properly as reported in literature (Mahanty, 2023).
- **Online model re-training to enable real-time learning:** Automated re-training is essential to empower the algorithm with “learning” capabilities. This feature exploits the fact that in parallel experiments the information generated by neighboring bioreactors is very useful for the running system.
- **A toolset to transfer learnings from past projects:** The software was able to describe perfusion runs with a new clone (Clone B), for which no previous data was available. This is possible because the GP Kernel can extract the existing similarities between the new clone and the known one. By this the existing information can be used to drastically reduce the number of experiments required in development.
- **Predictive process monitoring and notifications:** Despite the significant autonomy of the robotic system, auxiliary resources and external tasks (liquid containers, at-line analytics, sample handling for off-line analysis) still need to be performed by operators. In addition, unexpected malfunction in the bioreactors or liquid handlers need to be tackled by experts. Notifications that inform of an increasing probability that undesired events will take place (low dissolved oxygen, low glucose concentration, deviation on the pH set-point) allow acting on these events preventively.
- **Autonomous feed-back experimental operation minimizing the human in the loop:** One of the key advantages of robotics systems is operation 24/7. Processes that run in tightly defined conditions use process control to ensure robust operation, yet in development, neither a good process understanding (typically in formulated as a mechanistic mathematical model), nor a defined operating regime (as in manufacturing) are available. The software agent in charge of the operation of the parallel cultivation system is confronted with the challenging tasks to learn the new process behavior and simultaneously take actions that drive it to the desired targets (Nair et al., 2022). Furthermore, some targets set in the use case (e.g. cell viability) do not have a known input-output relationship, and it must be “learned” by the software during experimentation. It is hence vey relevant to demonstrate that the system was indeed able to find the process conditions that drive the process to the desired objective in all cases.

The above-described capabilities make this software solution a viable asset for process development, where high throughput systems are used for optimization of process conditions.

## 6. Conclusion

The introduction of autonomous experimental facilities (self-driving laboratories) in the pipeline of biopharma development is an essential step to accelerate the long and uncertain path for product to patient. The software solution presented here tackles important challenges by exploiting the potential of advanced parallel mini-bioreactor experimental systems. We demonstrate the capabilities of the developed software with three use cases, showing the added value of the implementation of machine learning tools in modern experimental systems. Furthermore, the user friendliness of the software and support offered to operators and scientists make this software an important step towards human centric industry 5.0.

An important challenge that remains to be solved in the future is modelling and feedback operation considering product quality. Currently, the main CQAs can only be measured offline leading to a significant time gap between the sampling and data availability (up to weeks). Process Analytical Technology (PAT) tools that enable online or at-line quantification of glycoforms, HCPs, aggregates, fragments will further boost bioprocess development and operation.

We expect the results to motivate further development in this area and a larger acceptance in the industry. The current gap between robotic capabilities and the autonomy of the devices needs to be urgently addressed in bioprocess development for the biotechnological and biopharmaceutical industries to match the high expectations set to digital biotechnology and bioindustry 5.0.

## Acknowledgments

We would like to thank the entire USP process design team (Biotech Development Center, Switzerland) for the support with the design and execution of the experiments in the lab. Specific thanks go to Raphaël Guillot and Anaïs Roulet for the feedbacks and interactions with the Datahow team. We also want to thank the Digital team, especially Alain Gillieron and Alexandre Gilet for their continuous support and agility. In addition, we do not forget our former colleagues Nicolas-Julian Hilbold, Stefania Caso and David Brühlmann who largely contributed to set the stage and kick-off this collaboration a few years back.

We also want to thank Daniel Stoeckel and Sonja Hatz (Merck KGaA, Darmstadt, Germany) for support with strategic decision and project management and the Merck 350 Research Grants funding for financing this innovation effort.

The discussion with the Digital Twin Team at Datahow significantly improved the software solution, specifically front end. We want to highlight Giulia Cantini for the implementation of the OPC connection, allowing the software to monitor the ongoing process.

We also thank Antonios Papaemmanouil and Thierry Prud’homme from Hochschule Luzern – Technik & Architektur for their support in the interface communication and control with the ambr software.

The writing of this manuscript was assisted by grammar checker and language correction AI tools specifically Trinka, Claude 3 Haiku, and GPT-3.5 Turbo. All authors have read, corrected and verified all information presented in this manuscript and Supplementary Information.

## Author’s Contribution

**Claudio Müller:** Writing – original draft, data preparation, visualization, modeling, software, conceptualization. **Thomas Vuillemin:** Writing – original draft, experimental operation, experimental design, conceptualization. **Chethana Janardhana Gadiyar:** Writing – original draft, review & editing. **Jean-Marc Bielser:** Writing – original draft, review & editing, project lead, conceptualization. **Jonathan Souquet:** Writing – review & editing, funding acquisition. **Alessandro Fagnani:** Writing – original draft, data preparation, modeling, software, conceptualization. **Michael Sokolov:** Writing – review & editing, modeling, funding acquisition. **Moritz von Stosch:** Writing – review & editing, modeling. **Fabian Feidl:** Writing – review & editing. **Alessandro Butté:** Writing – review & editing, modeling, funding acquisition. **Mariano Nicolas Cruz Bournazou:** Writing – original draft, supervision, project lead, conceptualization, funding acquisition.

## Data Availability

All data used in this study can be made available upon request.

## Appendix

### 1. Use case experiment case studies

**Figure A1.**
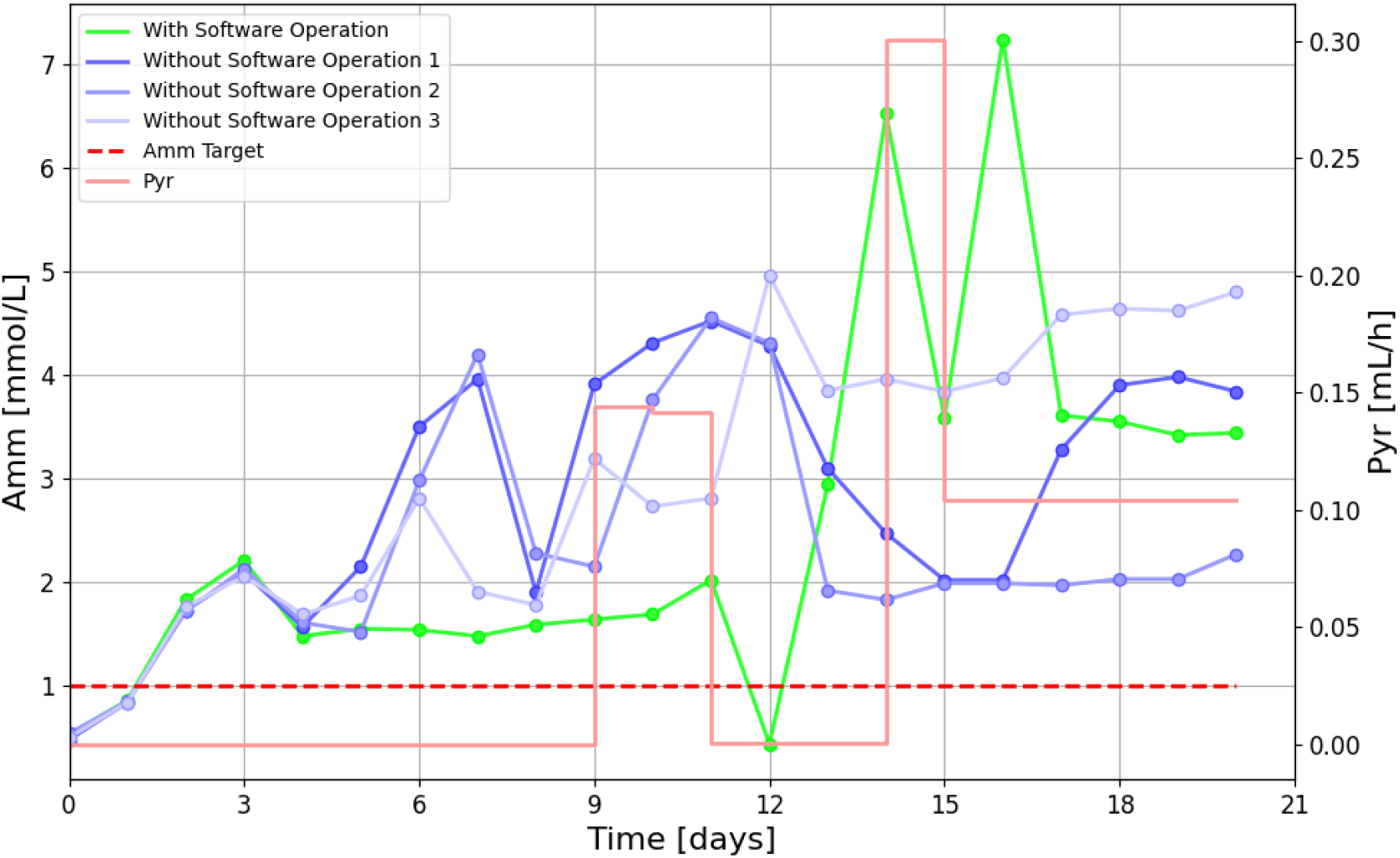
Condition 3 results: ammonium (Amm) use case with Amm target= 1 mM (red dashed line). The observed ammonium values of the bioreactor with software operation are shown in green, three bioreactors without software operation are shown in blue. The optimizer suggested pyruvate additions are plotted on secondary axis and is shown in light red. The model was originally not trained on data of this clone.

**Figure A2.**
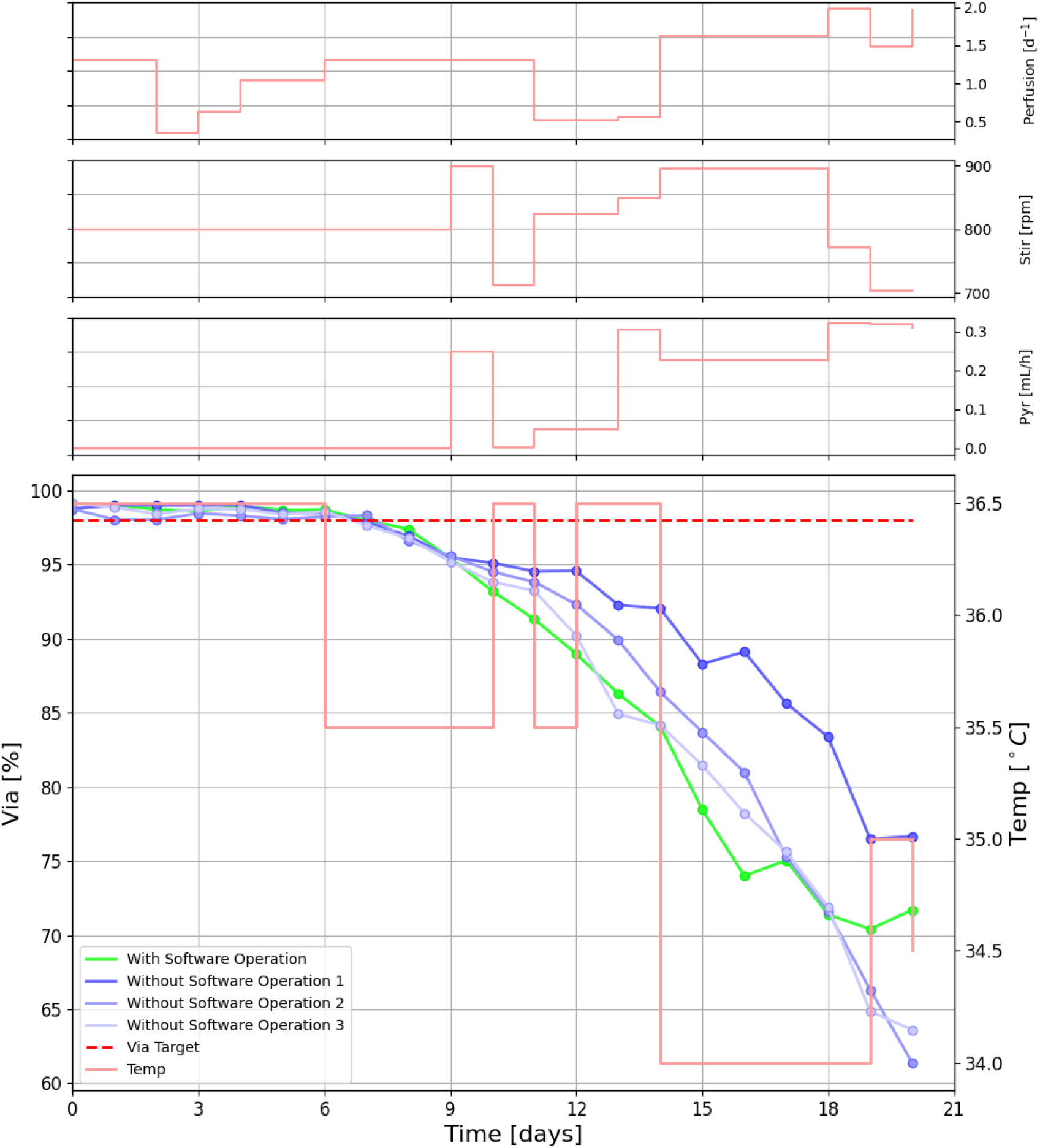
Condition 5 results: viability (Via) use case with Via Target = 98% (red dashed line). The observed viability values of the bioreactor with software operation are shown in green, three bioreactors without software operation are shown in blue. The optimizer suggested temperature rate is plotted on the secondary axis. All control parameter profiles are shown in light red.

**Figure A3.**
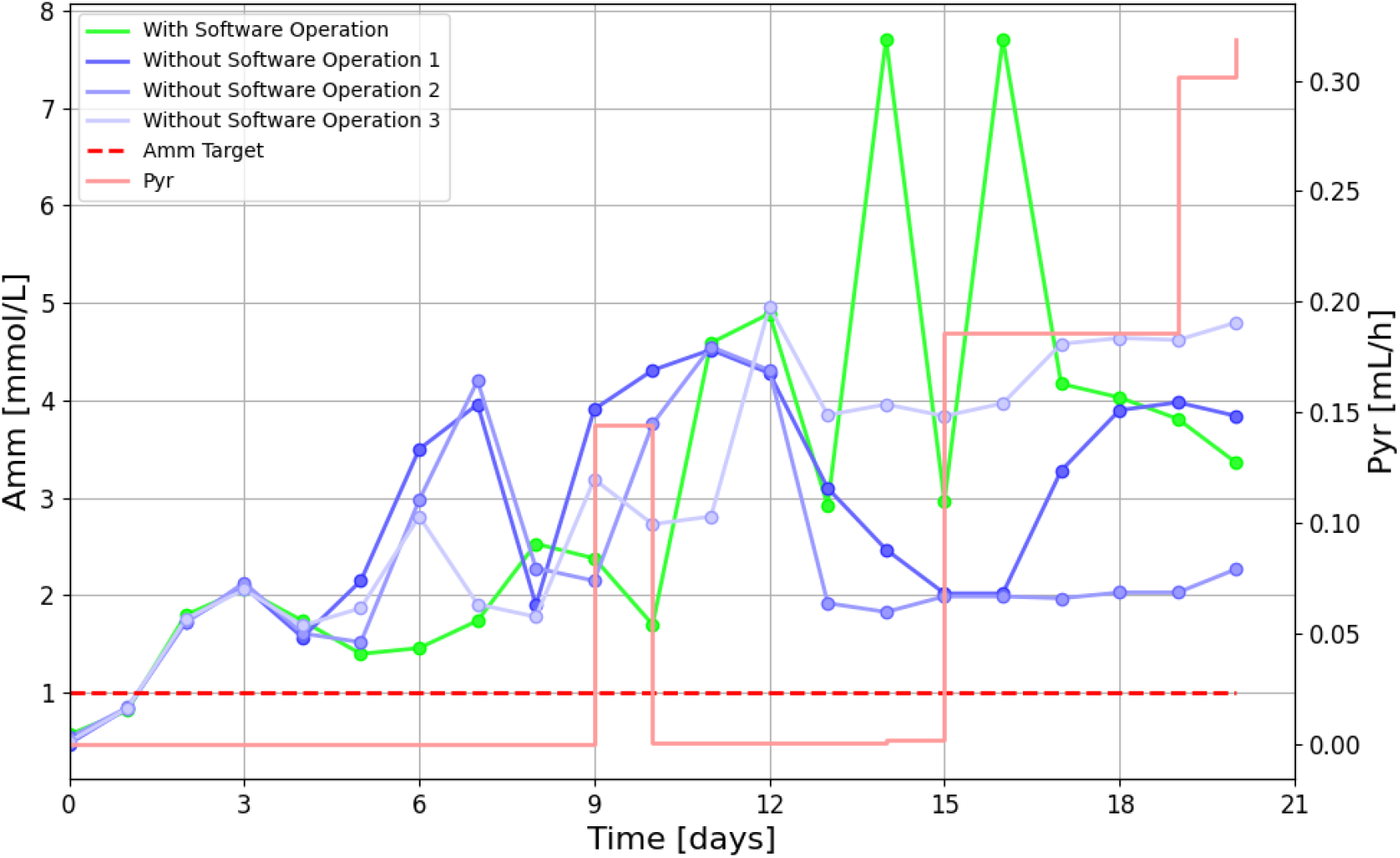
Condition 6 results: ammonium (Amm) use case with Amm target= 1 mM (red dashed line). The observed ammonium values of the bioreactor with software operation are shown in green, three bioreactors without software operation are shown in blue. The optimizer suggested pyruvate additions are plotted on secondary axis and is shown in light red. The model was originally not trained on data of this clone.

